## Supplementary information for "Signal Improved ultra-Fast Light-sheet Microscope (SIFT) for large tissue imaging"

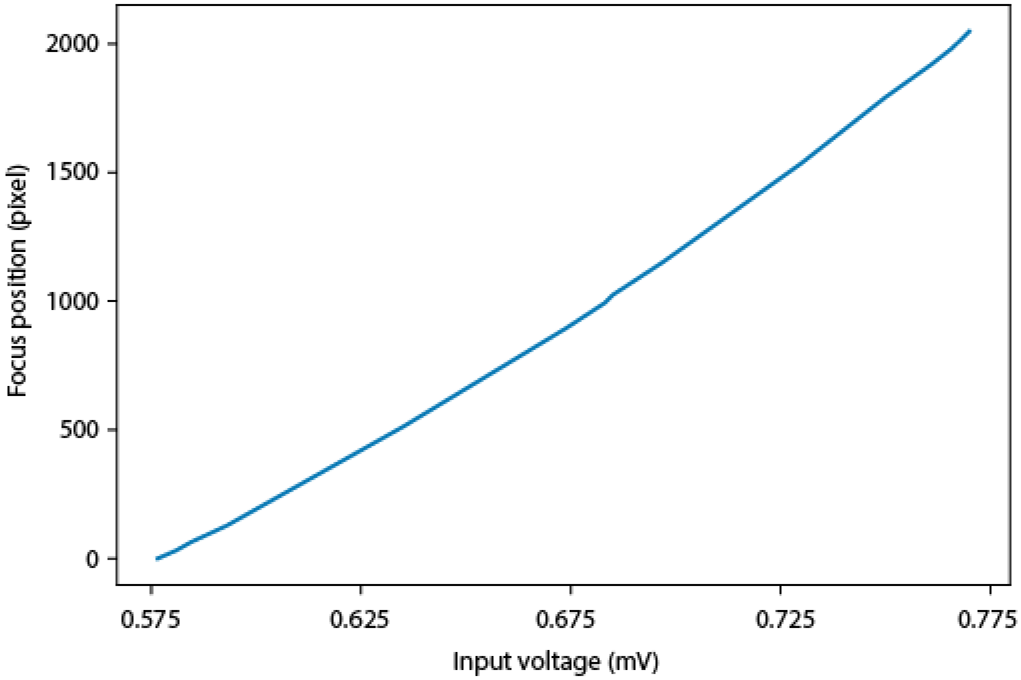


**Supplementary Fig. 1** | **LFA operating region.** Linear LFA movement with respect to the input voltage.

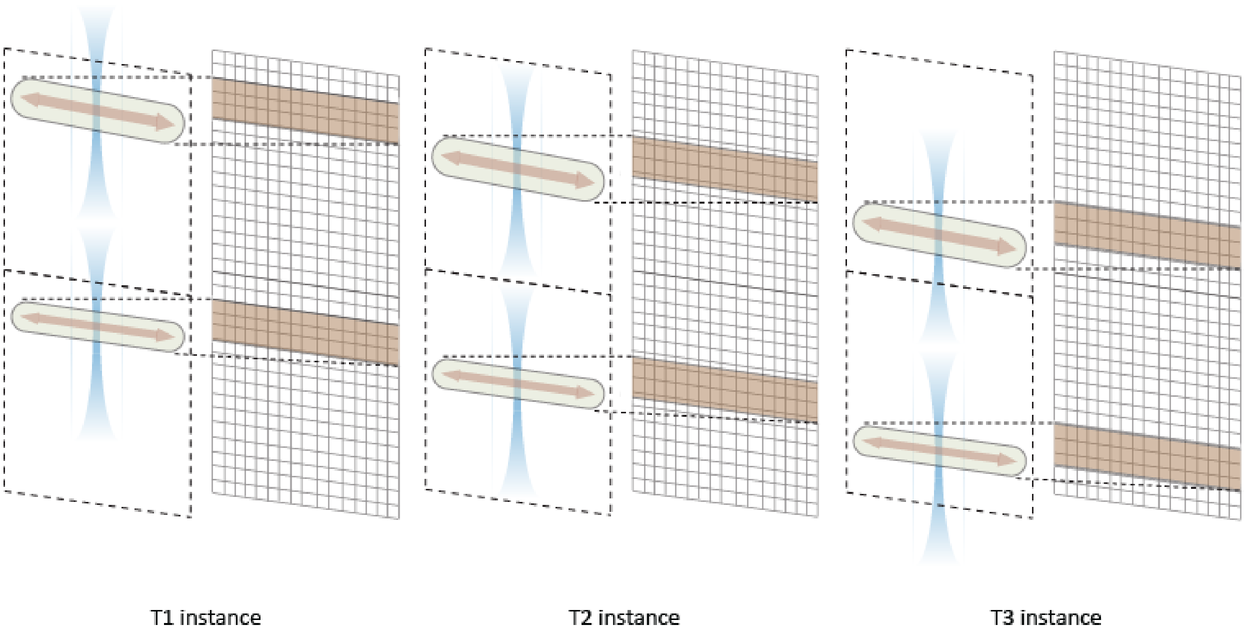


**Supplementary Fig. 2 | 2D focus scanning in SIFT.** Synchronous movement of dual 2D focus with the sCMOS camera rolling shutter, keeping a fixed separation between the two foci. The beam waists are captured by the camera chip.

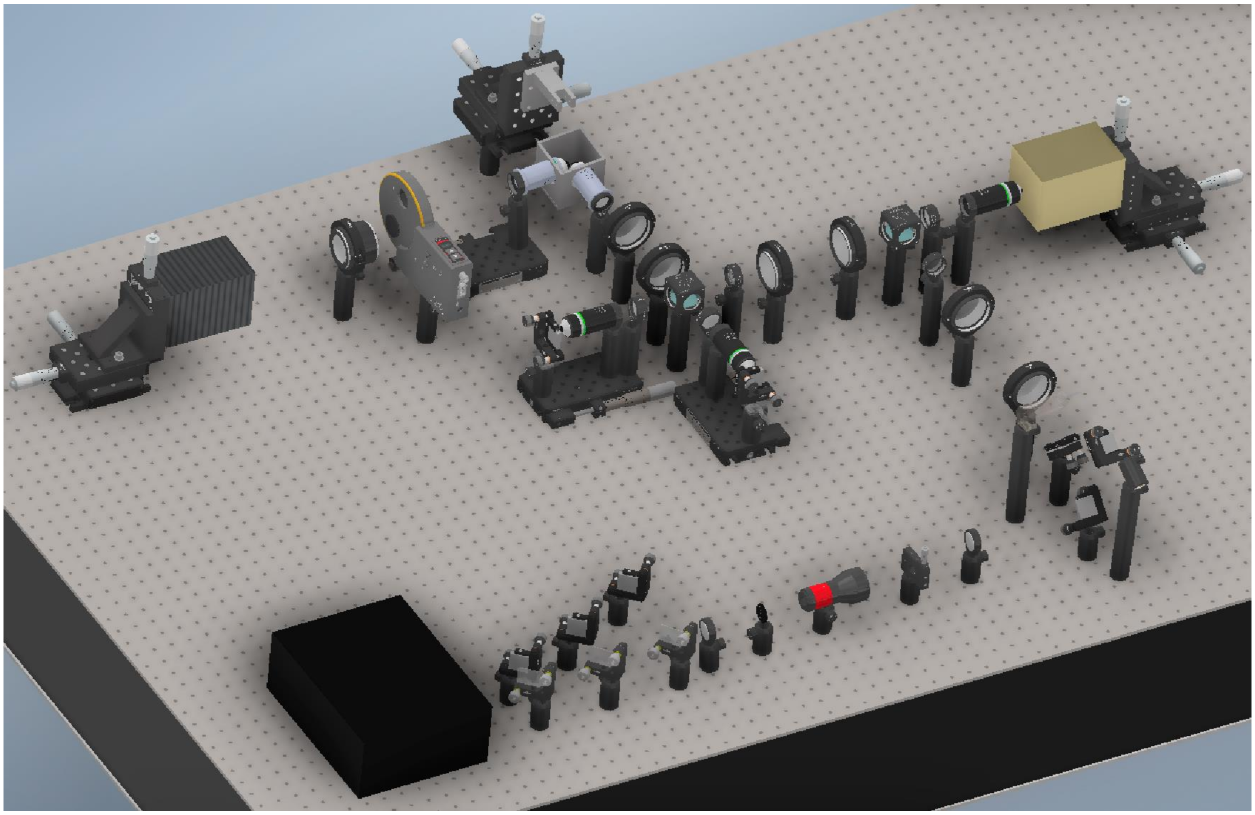

**Supplementary Fig. 3** | **Experimental implementation of SIFT.** 2D experimental implementation of SIFT. The necessary equipment list and the 3D simulated movie of SIFT implementation are delineated in **Supplementary Table 1** and **Supplementary Movie 1** respectively.

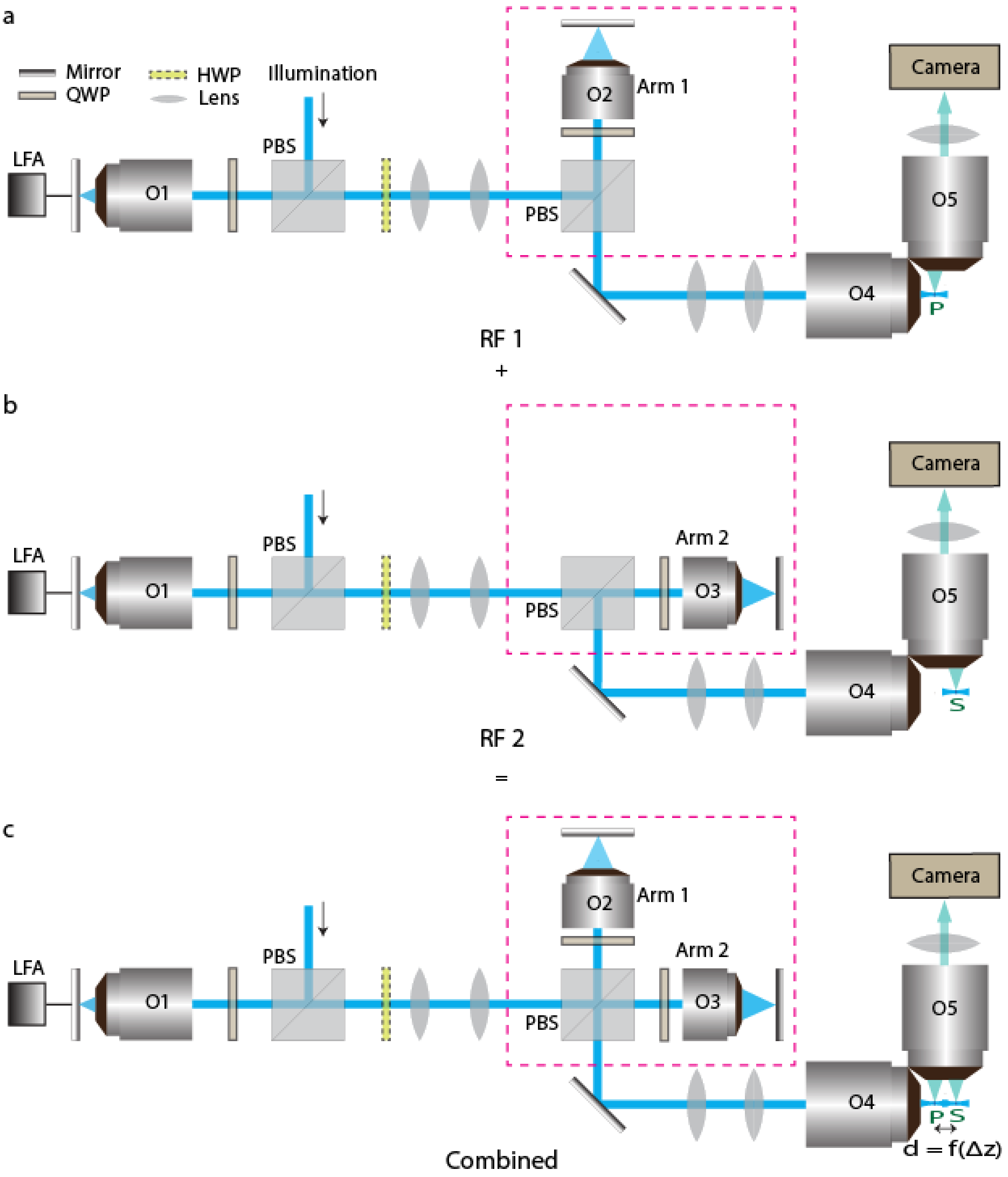


**Supplementary Fig. 4** | **Generation of two staggered light-sheet. a-b,** Two identical remote focusing (RF) path (RF1 and RF2). The RF1 includes O1, O2 and O4 where O1 and O2, O2 and O4 are pupil matched using the lens pair between them (**a**). Again, RF2 includes O1, O3 and O4 where O1 and O3, O3 and O4 are pupil matched (**b**). **c,** Combining RF1 and RF2 generates two LSs in the sample space. Different defocus parameter of the two path introduces a fixed distance between the two foci.

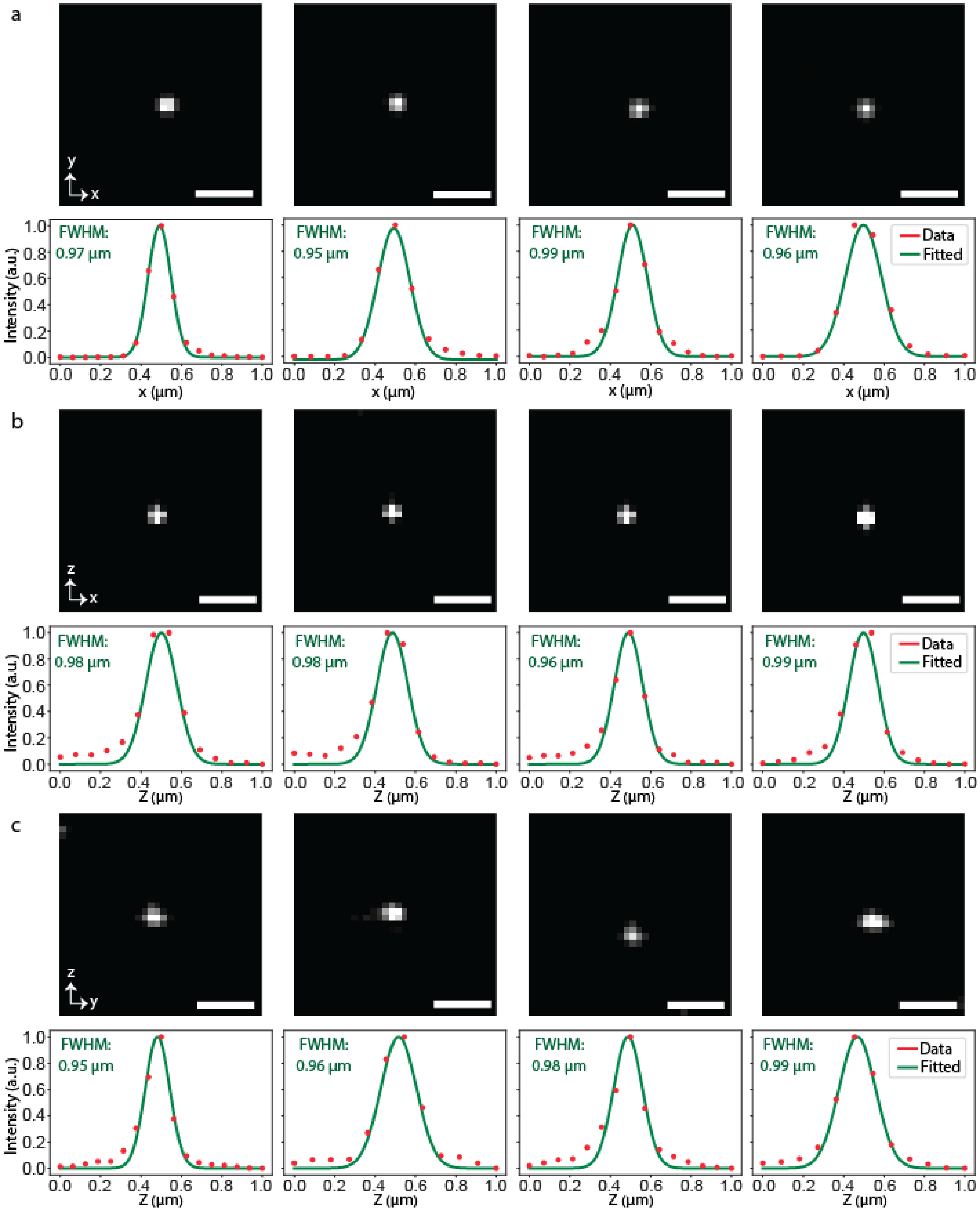


**Supplementary Fig. 5** | **Isotropic PSF. a-c,** SIFT enables imaging with isotropic resolution. 1st row shows four 500 nm fluorescent beads (imaged in water) and the 2nd row shows the FWHM of the PSF in XY (**a**), XZ (**b**) and YZ (**c**) dimensions. The isotropic imaging modality of SIFT provides the resolution of around 0.97 ± 0.05 µm in all dimensions. Scale bars, 3 µm (**a-c**).

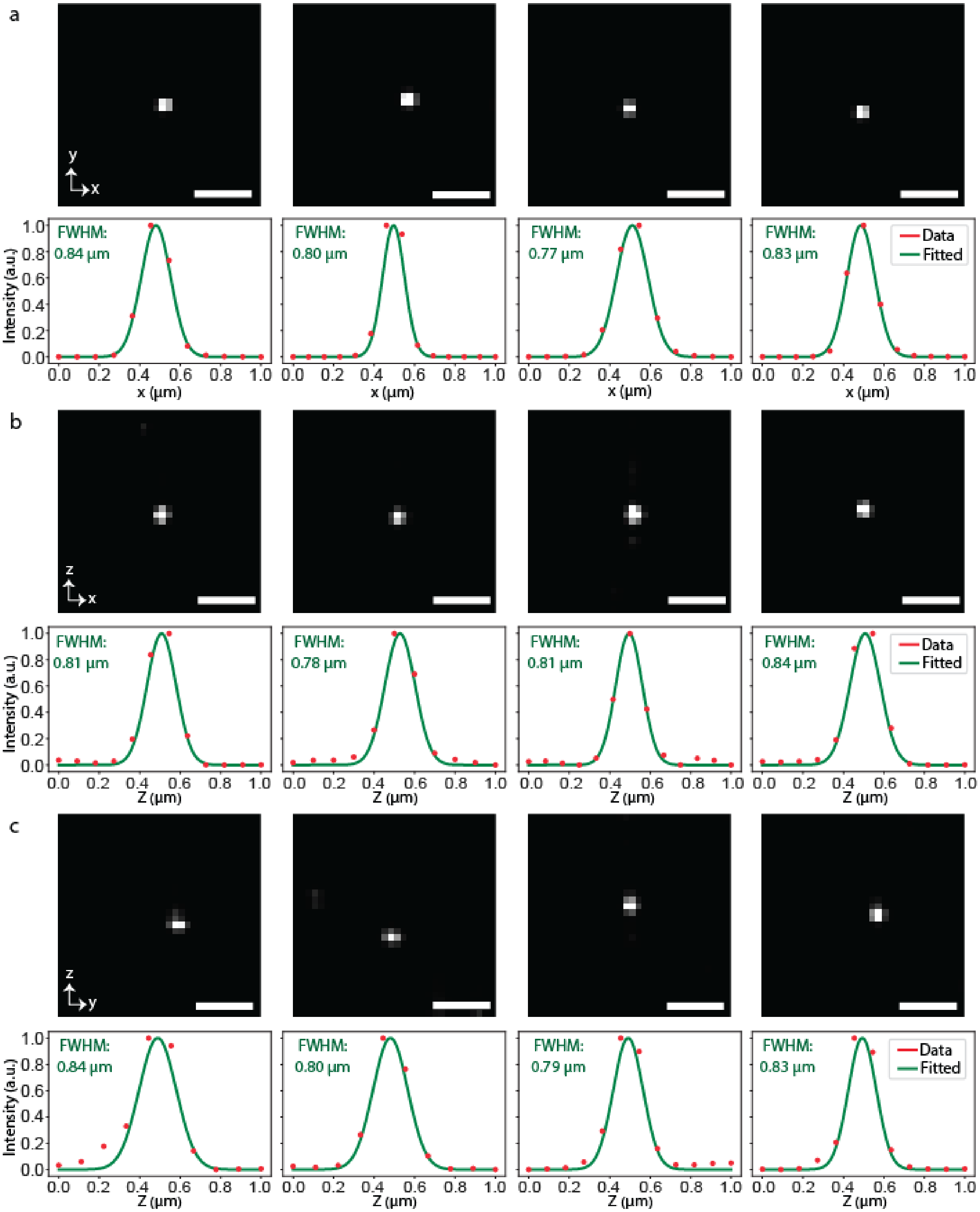


**Supplementary Fig. 6** | **Deconvolved isotropic PSF. a-c,** Resolution improvement following deconvolution. Several deconvolved (Richardson-Lucy) 500 nm fluorescent beads (imaged in water) are shown in the 1$^{st}$ row and the 2$^{nd}$ row shows the FWHM of the deconvolved PSF in XY (**a**), XZ (**b**) and YZ (**c**) dimensions. The isotropic imaging modality of SIFT provides the deconvolved resolution of around 0.80 ± 0.06 µm in all dimensions. Scale bars, 3 µm (**a-c**).

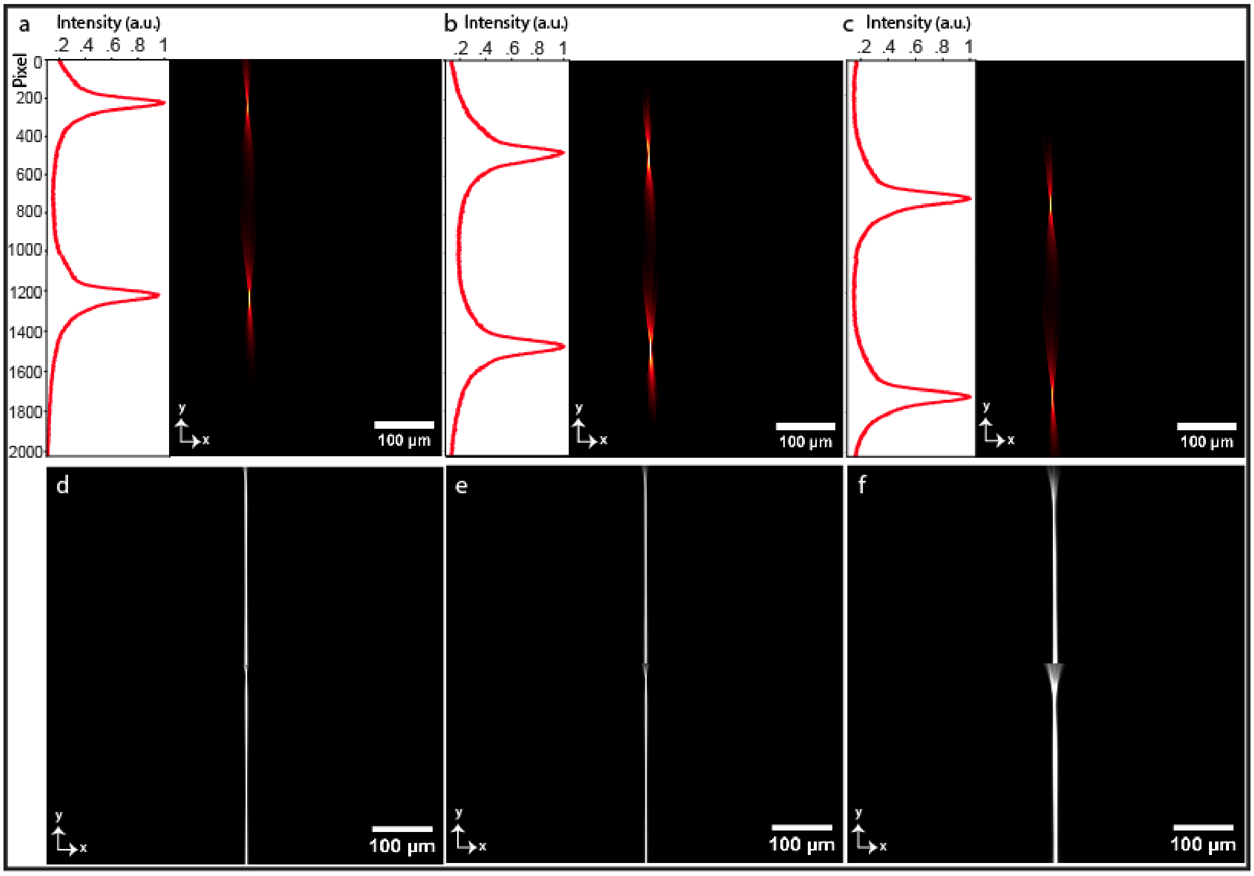


**Supplementary Fig. 7 | Synchronization of LS movement with camera rolling shutter. a-c,** Light-sheets along with the vertical intensity profile for the various positions of the camera FOV. The intensity profile ensures no out-of-focus light overlaps between the two LSs while travelling throughout the FOV. **d-f,** Scanning of 2D focus across the entire FOV for 25 ms (**d**), 20 ms (**e**) and 10 ms (**f**) of camera exposure time. The sharps line across the entire FOV depicts the tight synchronization between LSs and the rolling shutter which ensures uniform resolution across the entire FOV. It is also proven from **e** and **f** that with a small compromise of the FOV we may achieve the uniform resolution for even 20 ms or 10 ms of camera exposure time, five or ten-fold shorter than the exposure time of traditional ASLM.

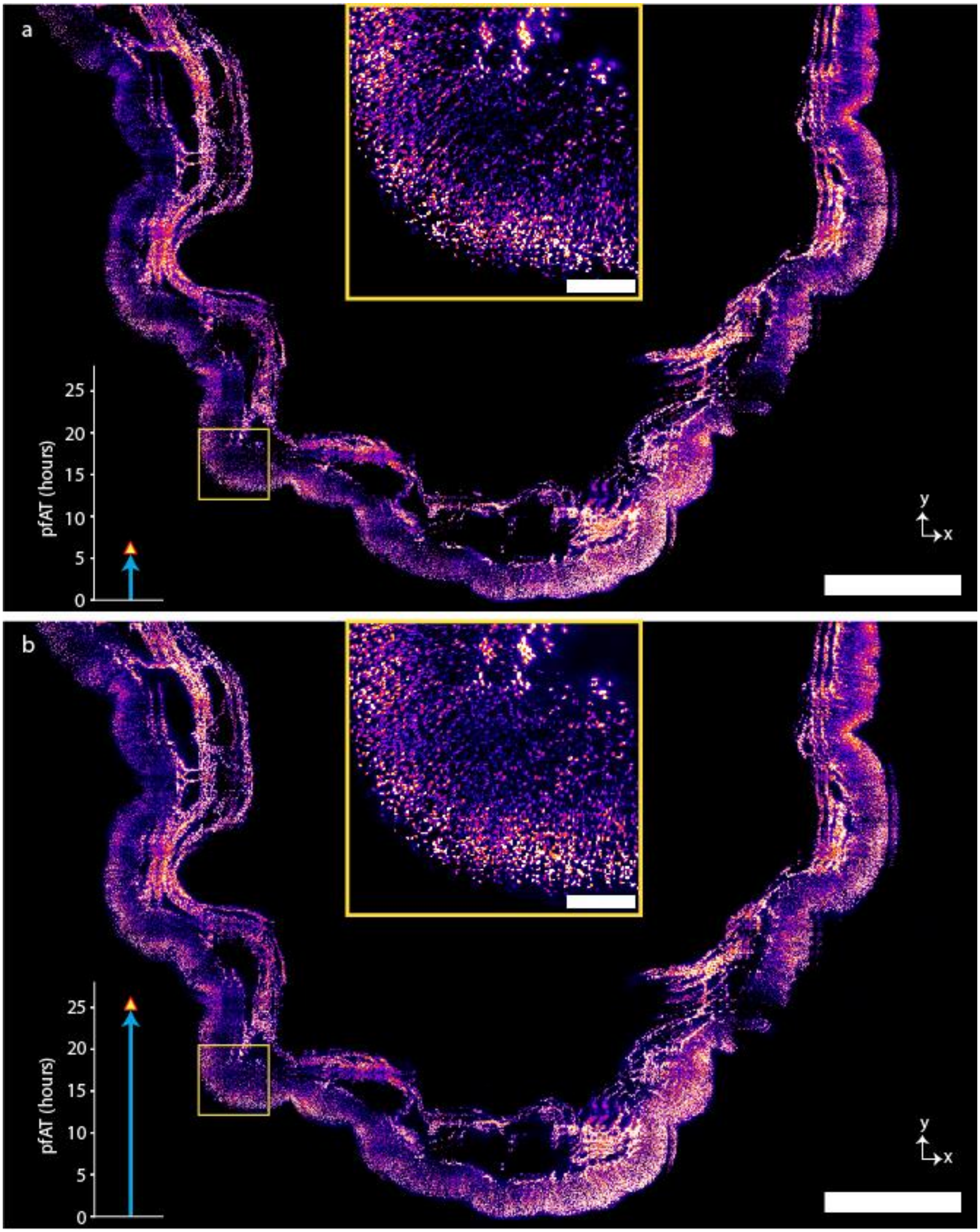


**Supplementary Fig. 8** | **Volumetric imaging of mouse stomach. a-b,** MIP of 100 slices of the whole stitched stomach, acquired by SIFT at 40 fps (**a**) and by traditional ASLM at 10 fps (**b**). Total pfAT is reflected by the bar plot of the respective images. Scale bars, 1 mm (**a,b**), 120 μm (inset of **a,b)**.

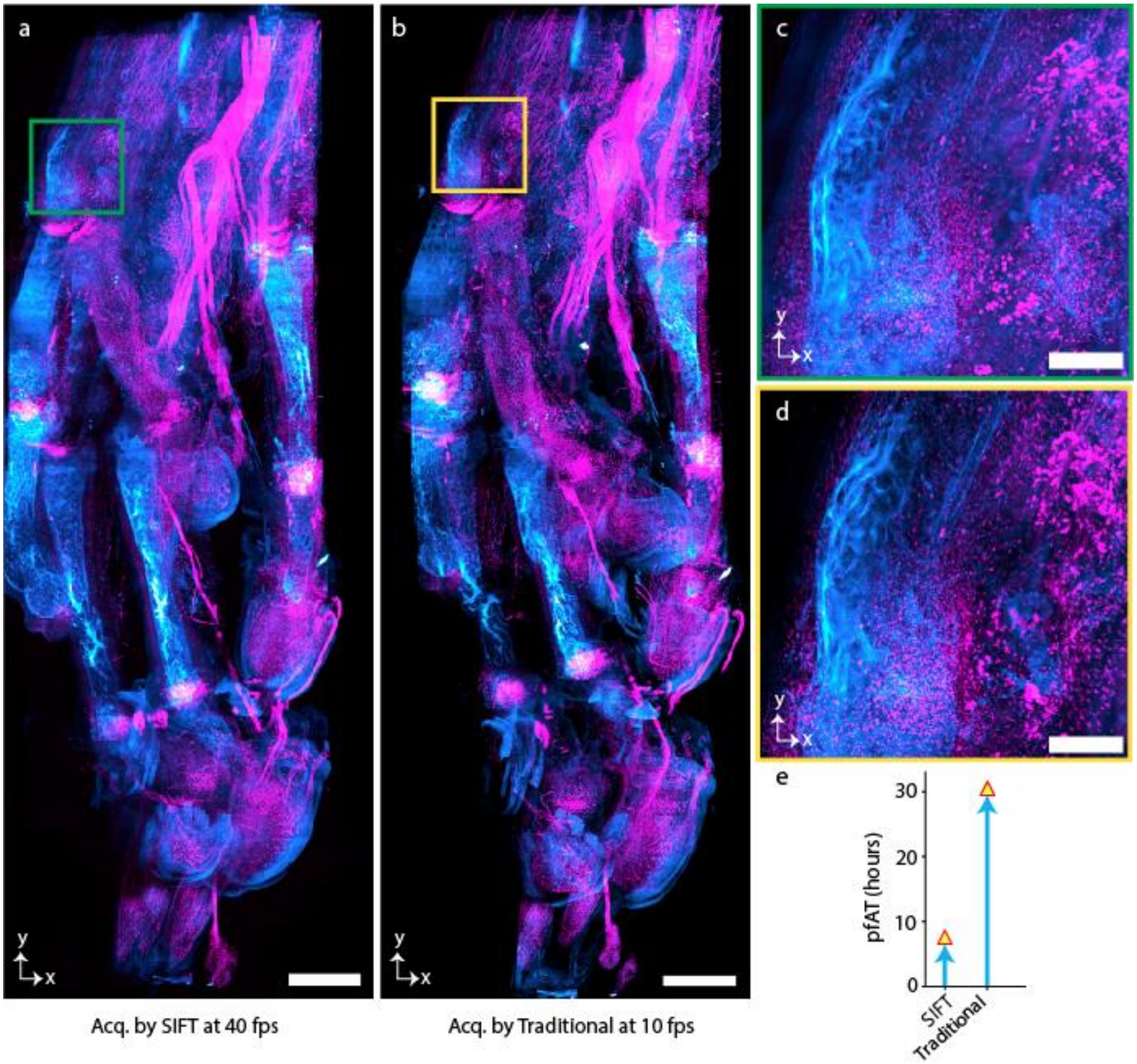


**Supplementary Fig. 9 | Volumetric imaging of mouse forepaw. a-b,** MIP of Wnt1-cre Scarlett flow dual channel mouse forepaw, acquired by SIFT at 40 fps (**a**) and by traditional ASLM at 10 fps (**b**). **c-d,** Higher magnification view of the randomly selected region from **a** and **b** respectively. **e,** Comparison of total pfAT to image the tissue using SIFT at 40 fps and traditional ASLM at 10 fps. Scale bars, 600 μm (**a, b**); 150 μm (**c,d**).

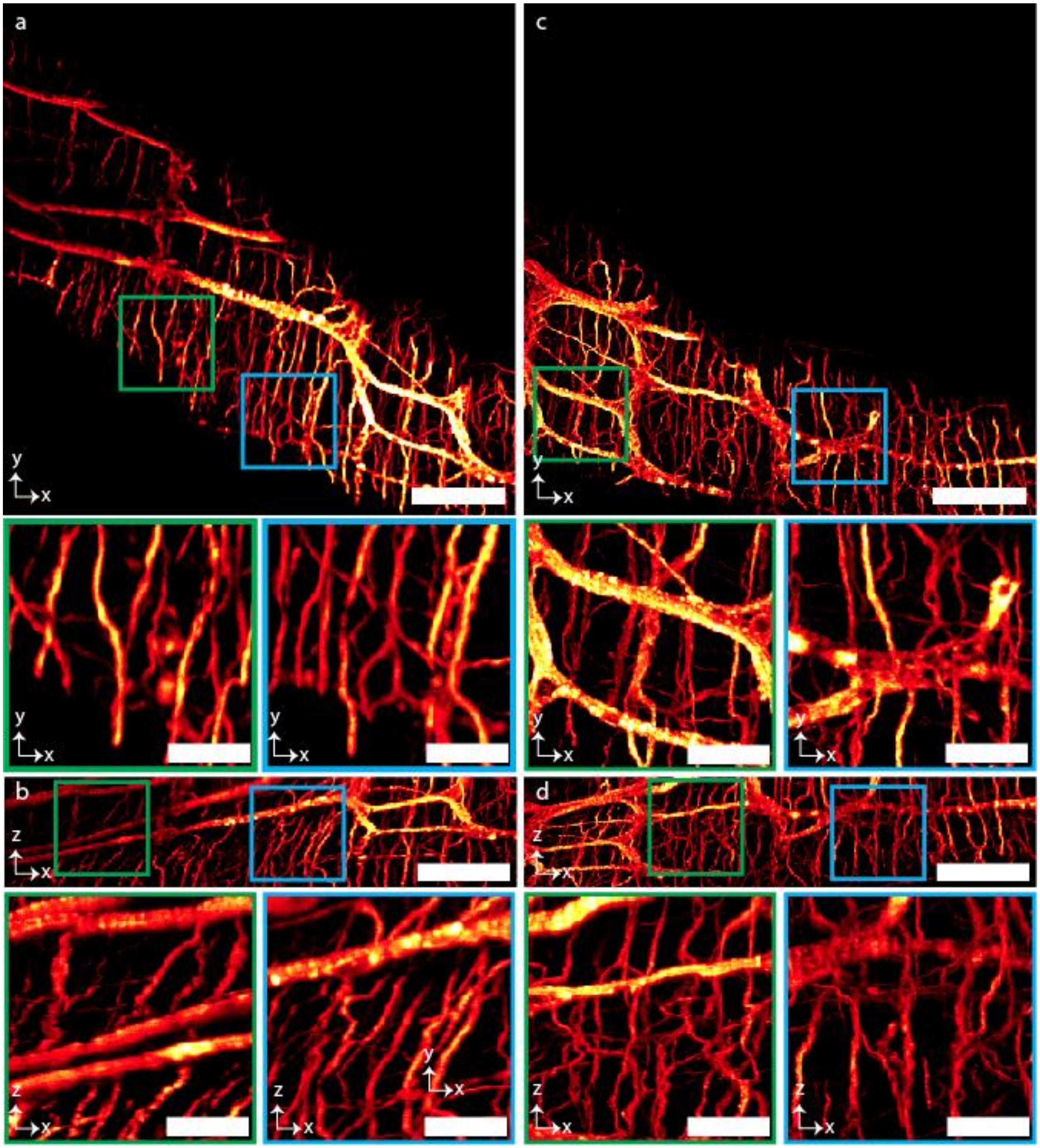


**Supplementary Fig. 10 | Volumetric imaging of mouse colon. a-b,** Lateral (**a**) and axial (**b**) view of the MIP of a tile of the mouse colon and the enlarged view of the randomly selected regions shown in square boxes of **a** and **b. c-d,** Lateral (**c**) and axial (**d**) view of the MIP of another tile of the mouse colon and the enlarged view of the randomly selected regions shown in square boxes of **c** and **d**. Scale bars, 120 μm (**a-d**), enlarged view 40 μm.

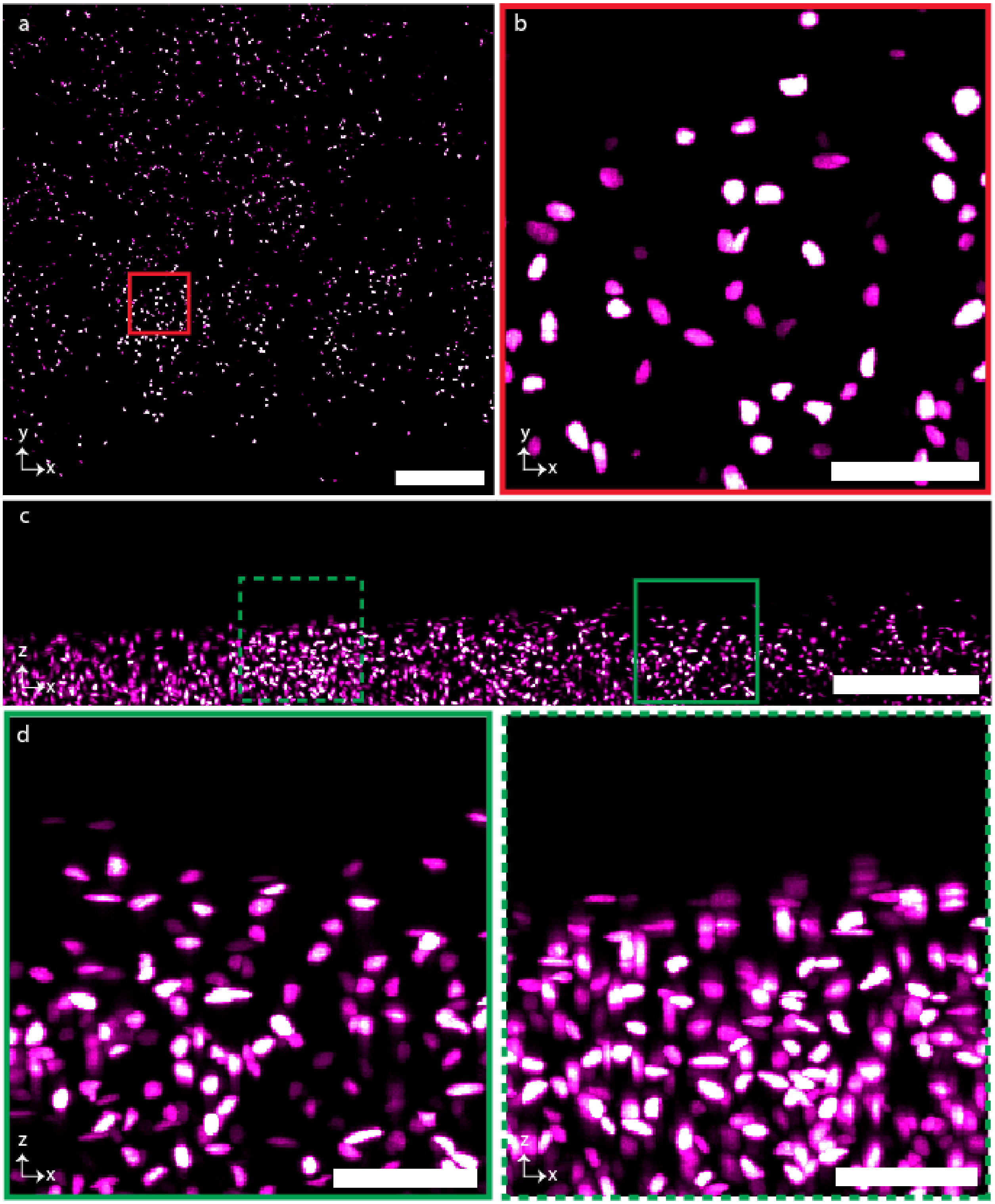


**Supplementary Fig. 11a | Representative tile of mouse stomach. a,** MIP of the lateral view of a single tile among the 2,171 tiles of mouse stomach (shown stitched stomach in **Supplementary Fig. 8** and **Supplementary Movie 3**). **b,** Higher magnification view of a randomly selected region from **a**. **c,** MIP of the axial view of the same tile. **d,** Higher magnification view of the selected regions of **c**. Scale bars, 120 µm (**a**); 45 µm (**b,d**); 100 µm (**c**).

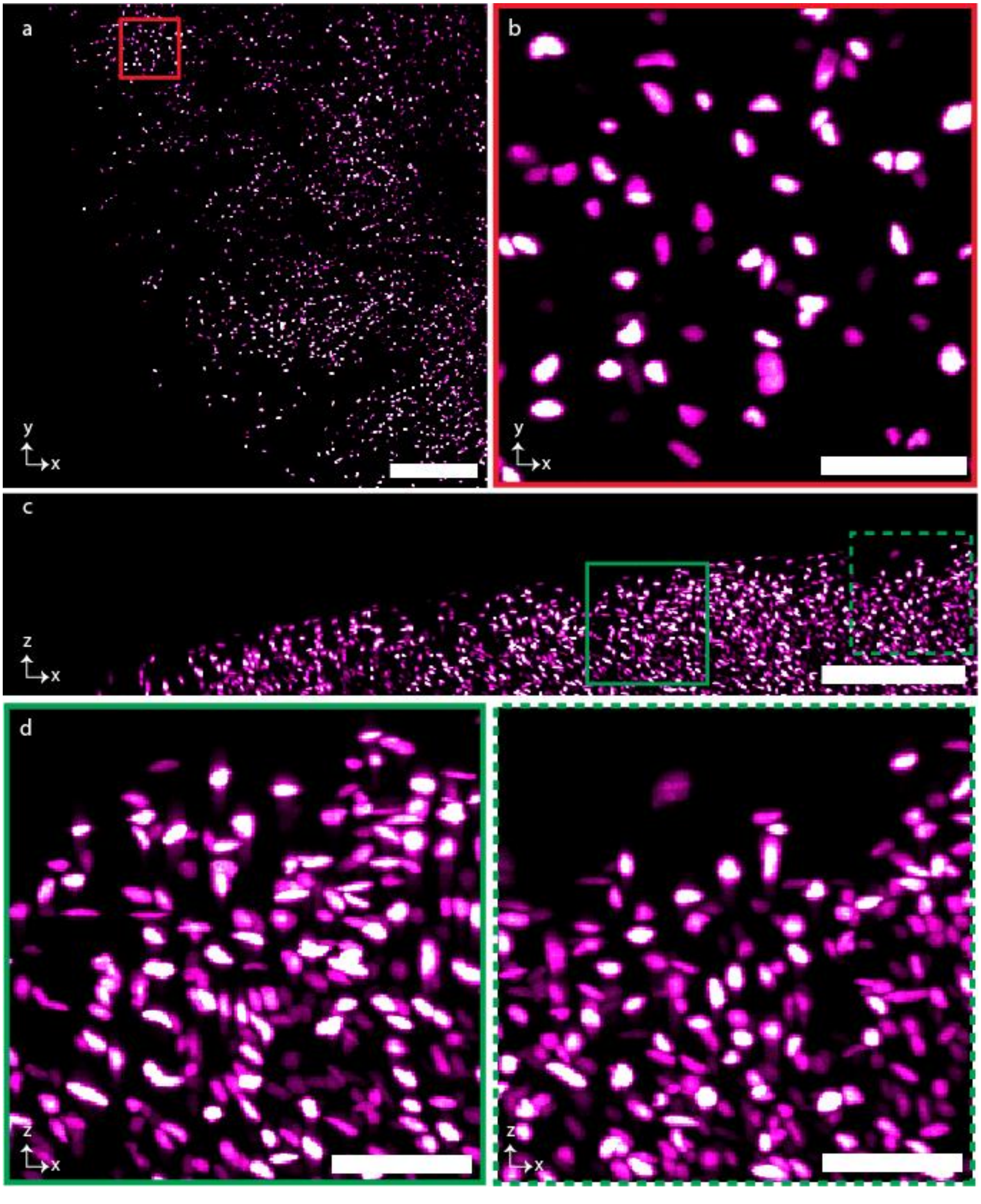


**Supplementary Fig. 11b | Representative 2nd tile of mouse stomach. a,** MIP of the lateral view of a single tile among the 2,171 tiles of mouse stomach (shown stitched stomach in **Supplementary Fig. 8** and **Supplementary Movie 3**). **b,** Higher magnification view of a randomly selected region from **a**. **c,** MIP of the axial view of the same tile. **d,** Higher magnification view of the selected regions of **c**. Scale bars, 120 μm (**a**); 45 μm (**b,d**)**;** 100 μm (**c**).

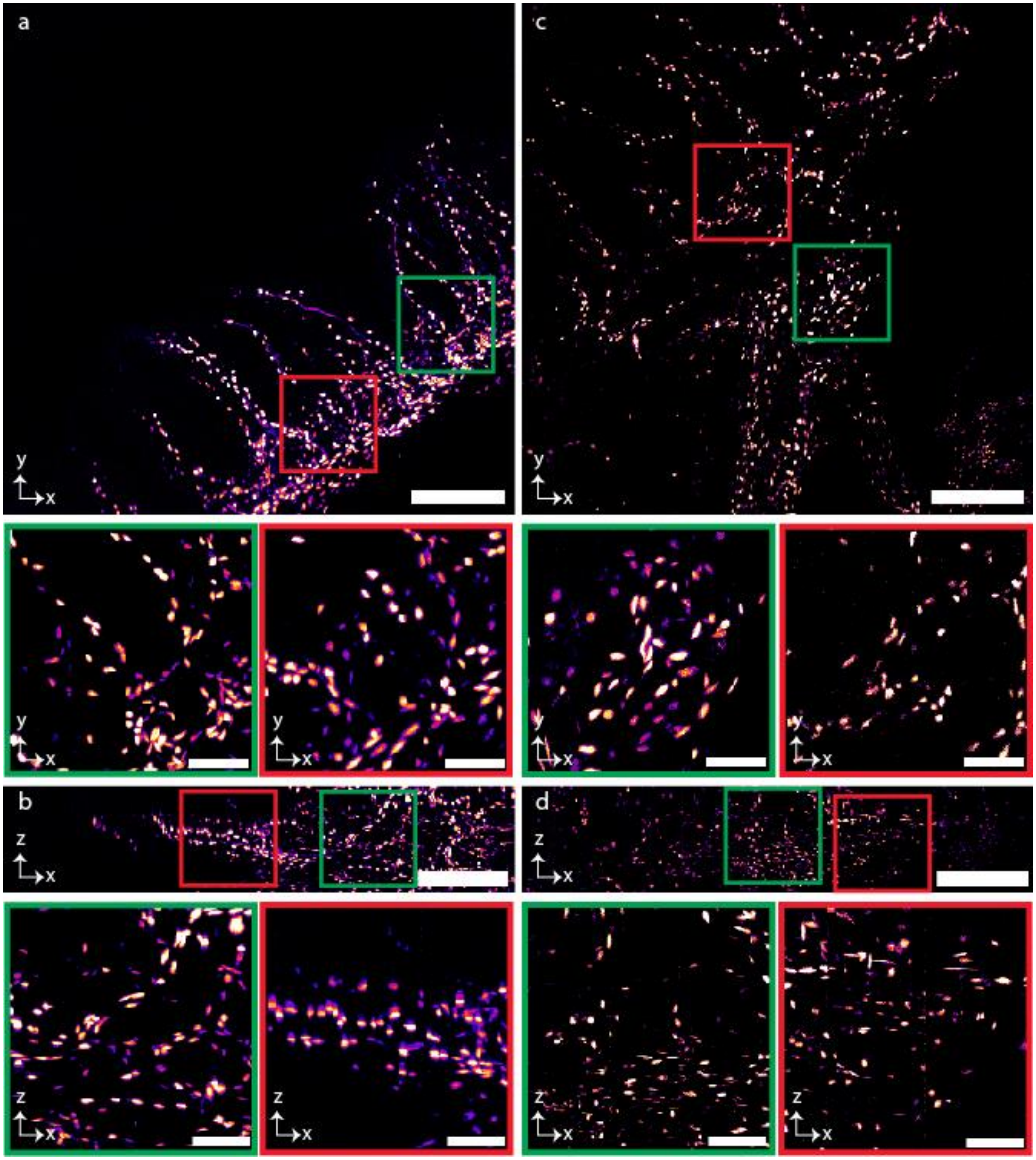


**Supplementary Fig. 12 | Image of different tiles of mouse gut. a-b,** Lateral (**a**) and axial (**b**) view of the MIP of a tile of the mouse gut and the enlarged view of the randomly selected regions shown in square boxes of **a** and **b.** (shown stitched gut in **Extended Data Fig. 3** and **Supplementary Movie 8**). **c-d,** Lateral (**c**) and axial (**d**) view of the MIP of another tile of the mouse gut and the enlarged view of the randomly selected regions shown in square boxes of **c** and **d**. Scale bars, 120 μm (**a-d**), 25 μm (enlarged view of **a-d**).

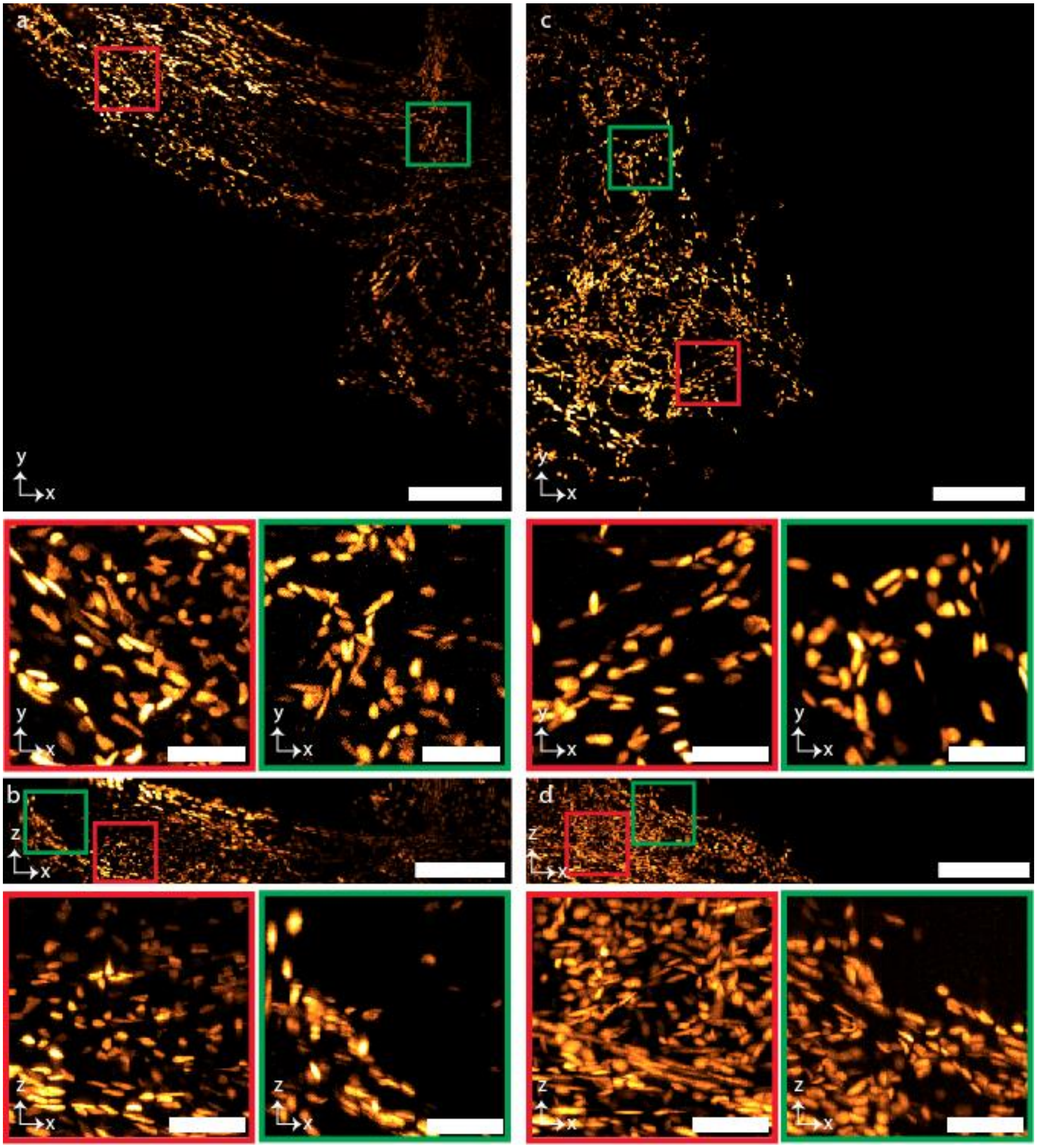


**Supplementary Fig. 13 | Image of different tiles of mouse hindpaw. a-b,** Lateral (**a**) and axial (**b**) view of the MIP of a tile of the mouse forepaw and the enlarged view of the randomly selected regions shown in square boxes of **a** and **b. c-d,** Lateral (**c**) and axial (**d**) view of the MIP of another tile of the mouse forepaw and the enlarged view of the randomly selected regions shown in square boxes of **c** and **d**. Scale bars, 120 μm (**a-d**), 25 μm (enlarged view of **a-d**).

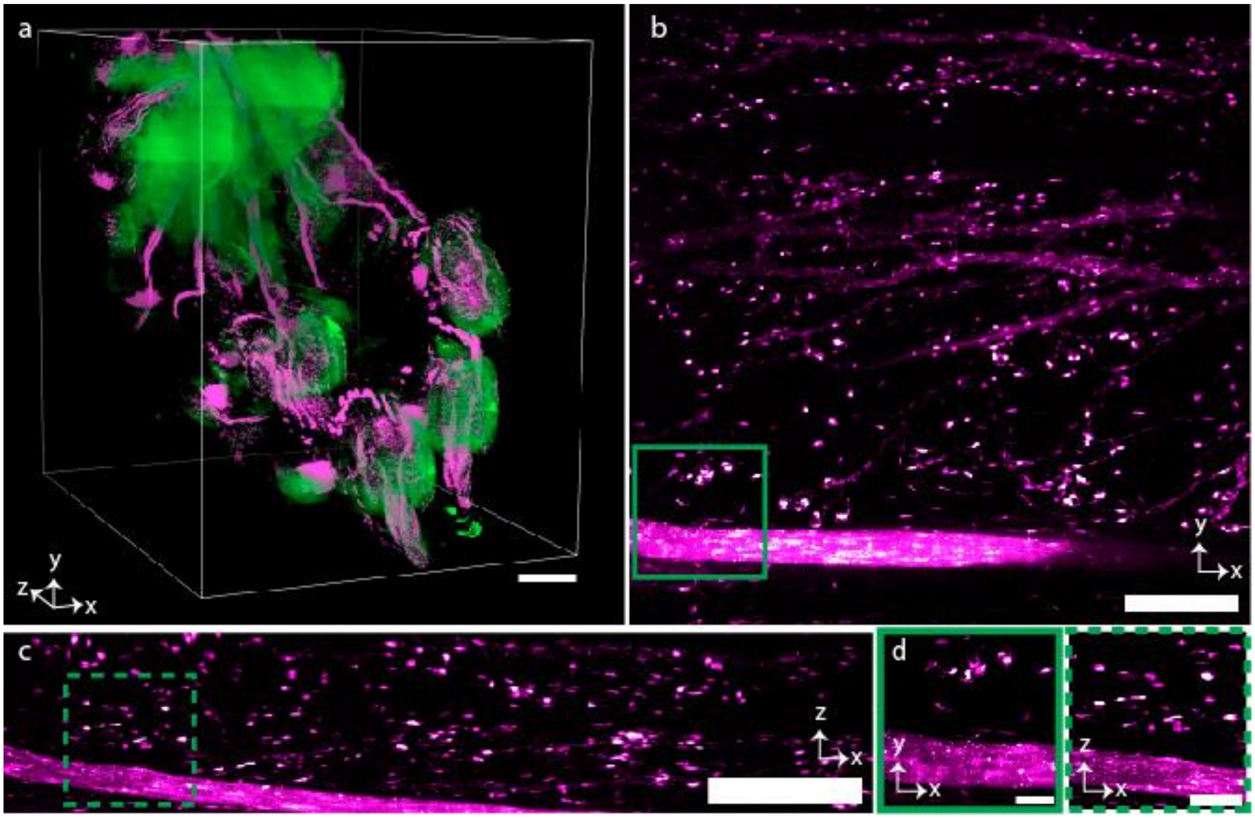


**Supplementary Fig. 14 | Volumetric imaging of mouse forepaw. a,** 3D rendered view of dual channel mouse forepaw (Fig. 3 and **Supplementary Movie 2**) cleared by PEGASOS protocol and imaged by SIFT at 40 fps. The image volume is 4.2 x 3.3 x 5.5 $mm^3$. **b-c,** MIP of the lateral (**b**) and axial (**c**) view of one image tile. **d,** Enlarged view of the selected regions from **b** and **c**, marked by green square boxes. Scale bars, 500 μm (**a**); 120 μm (**b,c**); 30 μm (**d**).

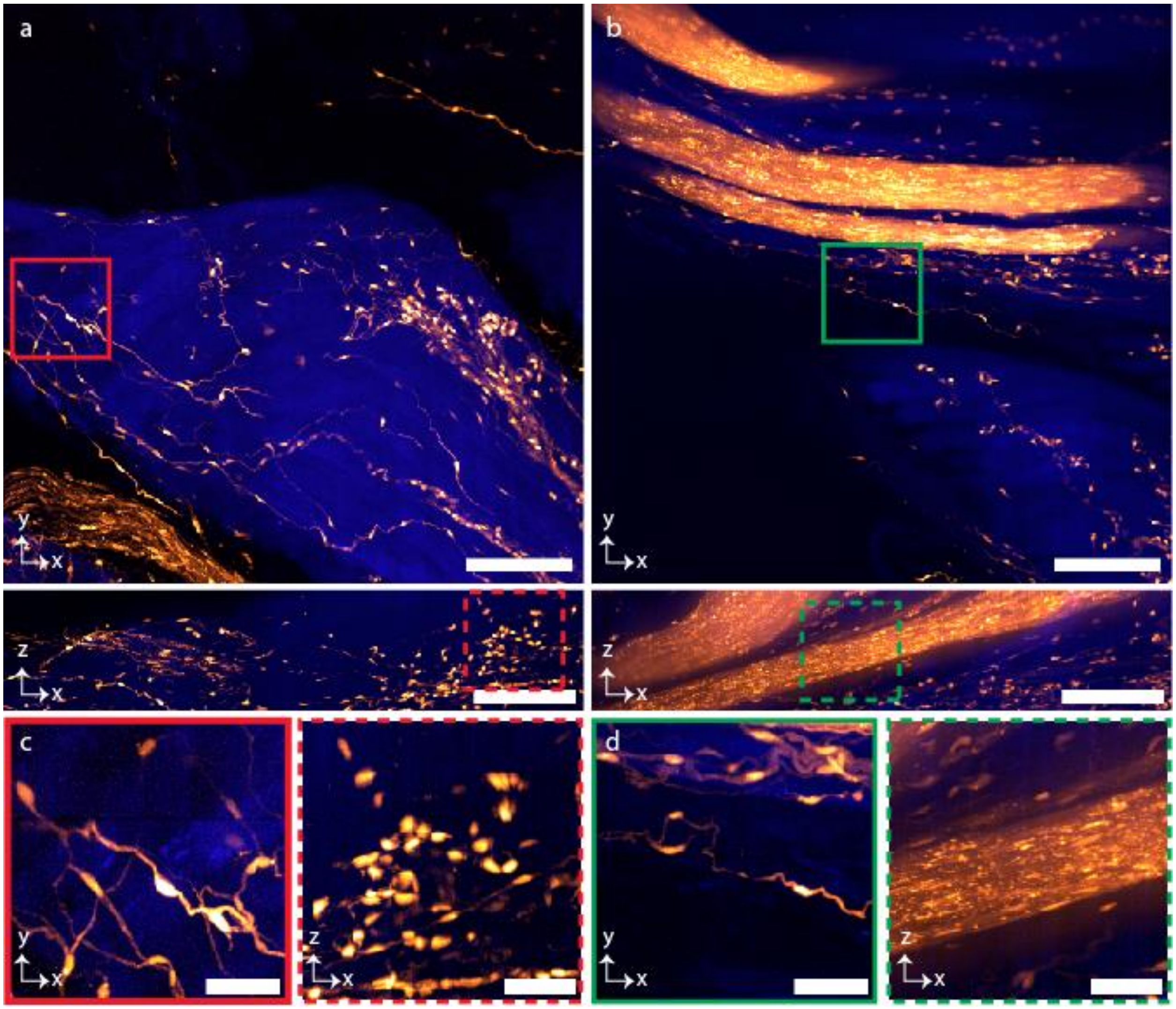


**Supplementary Fig. 15 | Image of different tiles of mouse forepaw. a-b,** Lateral and axial view of the MIP of a tile of the mouse forepaw for two random tiles (whole stitched dual channel forepaw is shown in **Fig. 3**). **c-d,** Higher magnification view of randomly selected regions from **a** (**c**) and **b** (**d**). Scale bars, 120 μm (**a-b**), 30 μm (**c-d**).

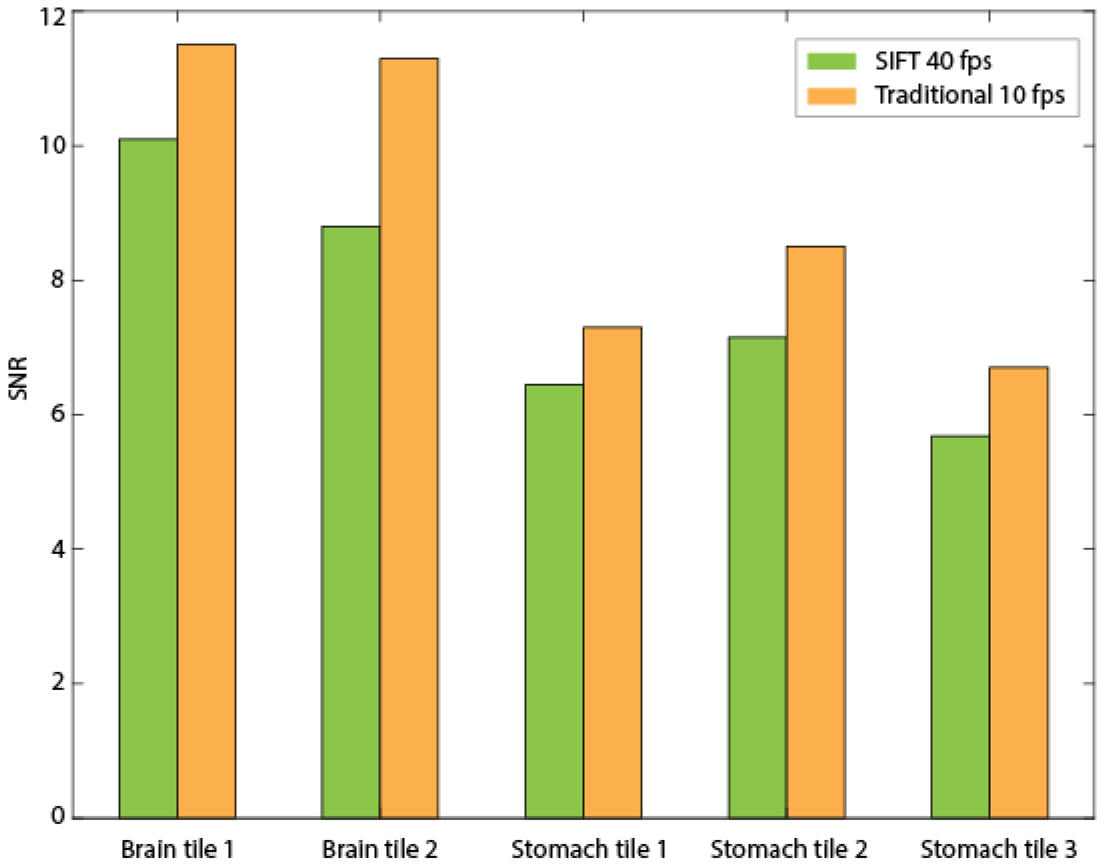


**Supplementary Fig. 16** | **Comparison of SNR.** Statistical data for signal to noise ratio (SNR), computed for randomly selected five tiles taken from two different specimens acquired by SIFT at 40 fps and traditional ASLM at 10 fps.

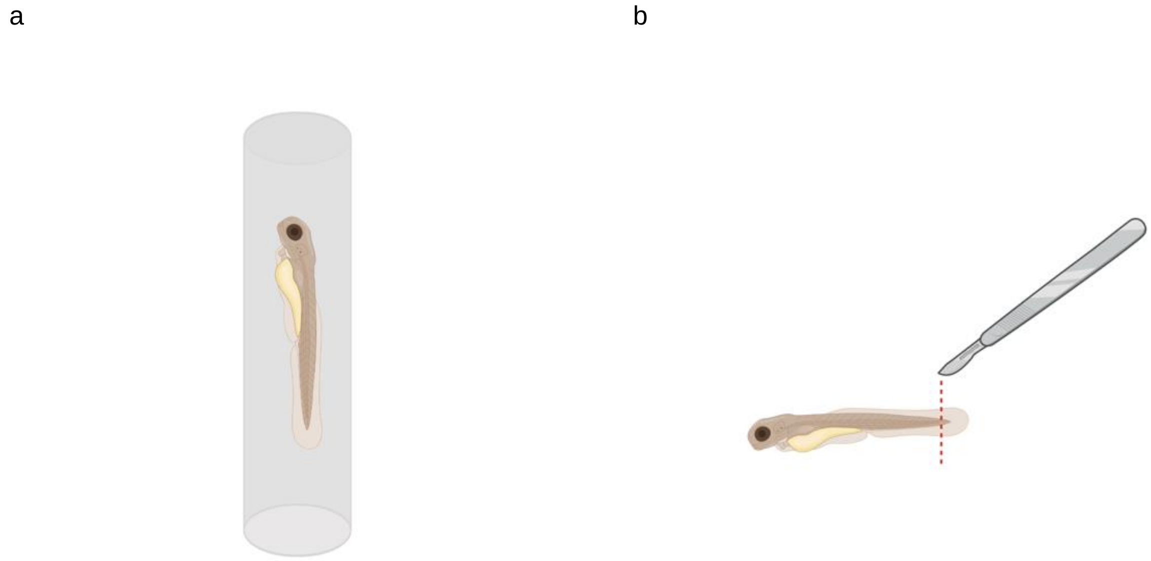


**Supplementary Fig. 17 | Imaging of live zebrafish. a,** A live zebrafish larvae, anesthetized, embedded in agarose and then mounted in a FEP tube. **b,** Methods for zebrafish tail cut. The larvae were anesthetized in Tricaine (160 mg/L) and tails were cut with a sharp scalpel. Fish were then placed in low melt agarose at 42˚C before being sucked into a capillary tube made of FEP (ZEUS Virtual item: 0000183678).

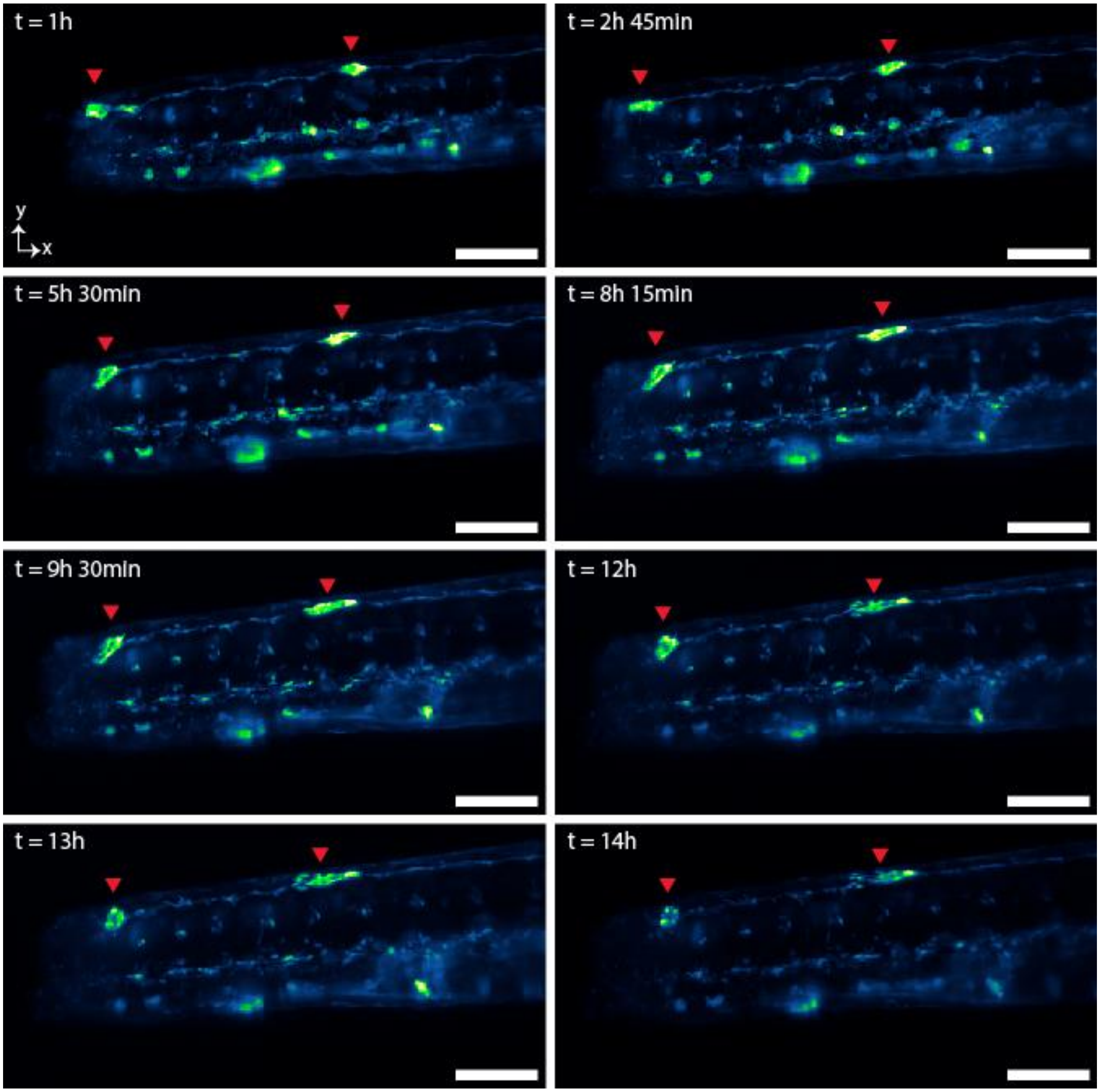


**Supplementary Fig. 18 | Additional z-depth of Zebrafish tail following injury.** As seen in Fig. 4, ROI as marked by the red box shown in Fig. **4a**, and images are of the same animal as seen in Fig. **4b** (different z-depth). Red arrows indicate regions of egfp expression with a migratory behavior or change in morphology. Scale bar, 100 μm.

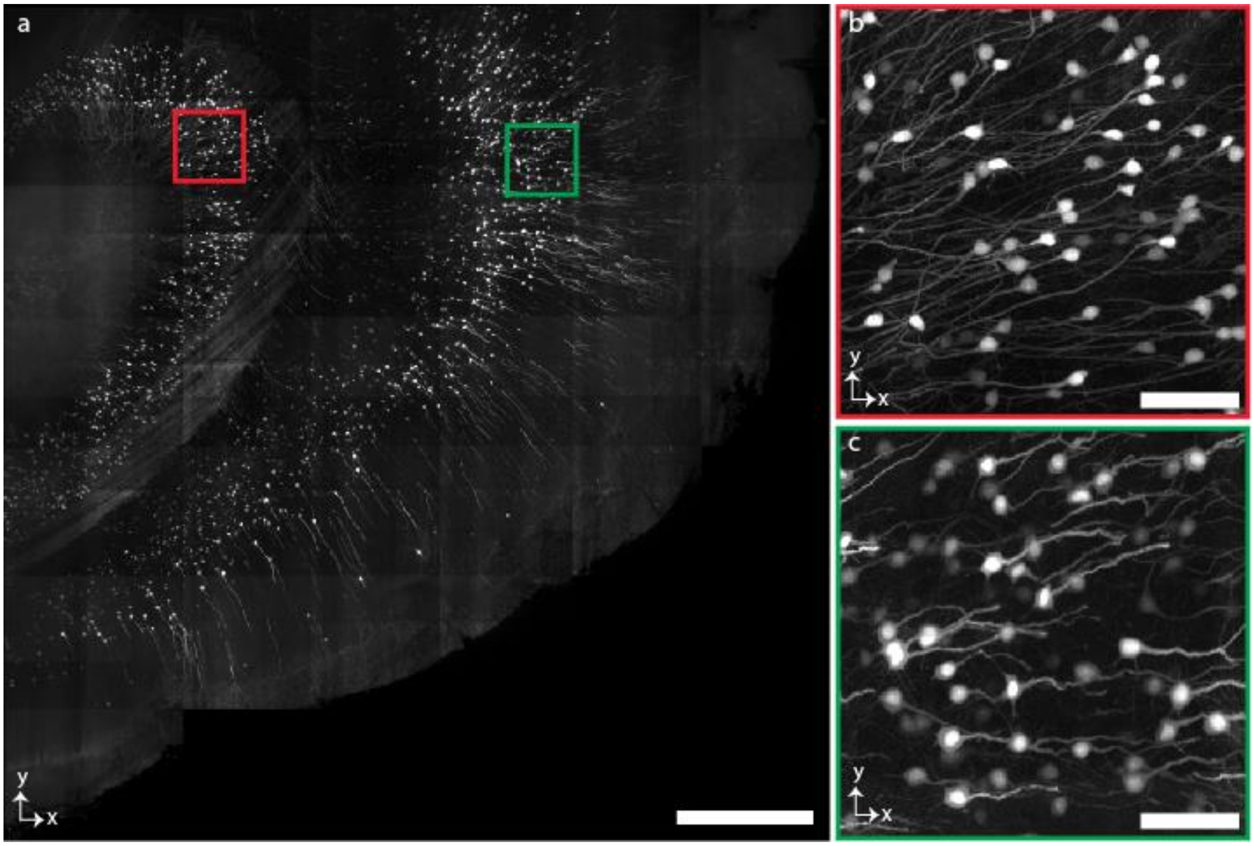


**Supplementary Fig. 19 | Image of Thy-1 GFP mouse brain. a,** MIP of a part of Thy-1 GFP neuronal mouse brain. **c-b,** Higher magnification view of randomly selected regions corresponding to the red (**b**) and green (**c**) square boxes of **a**. Scale bars, 500 μm (**a**), 60 μm (**b-c**).

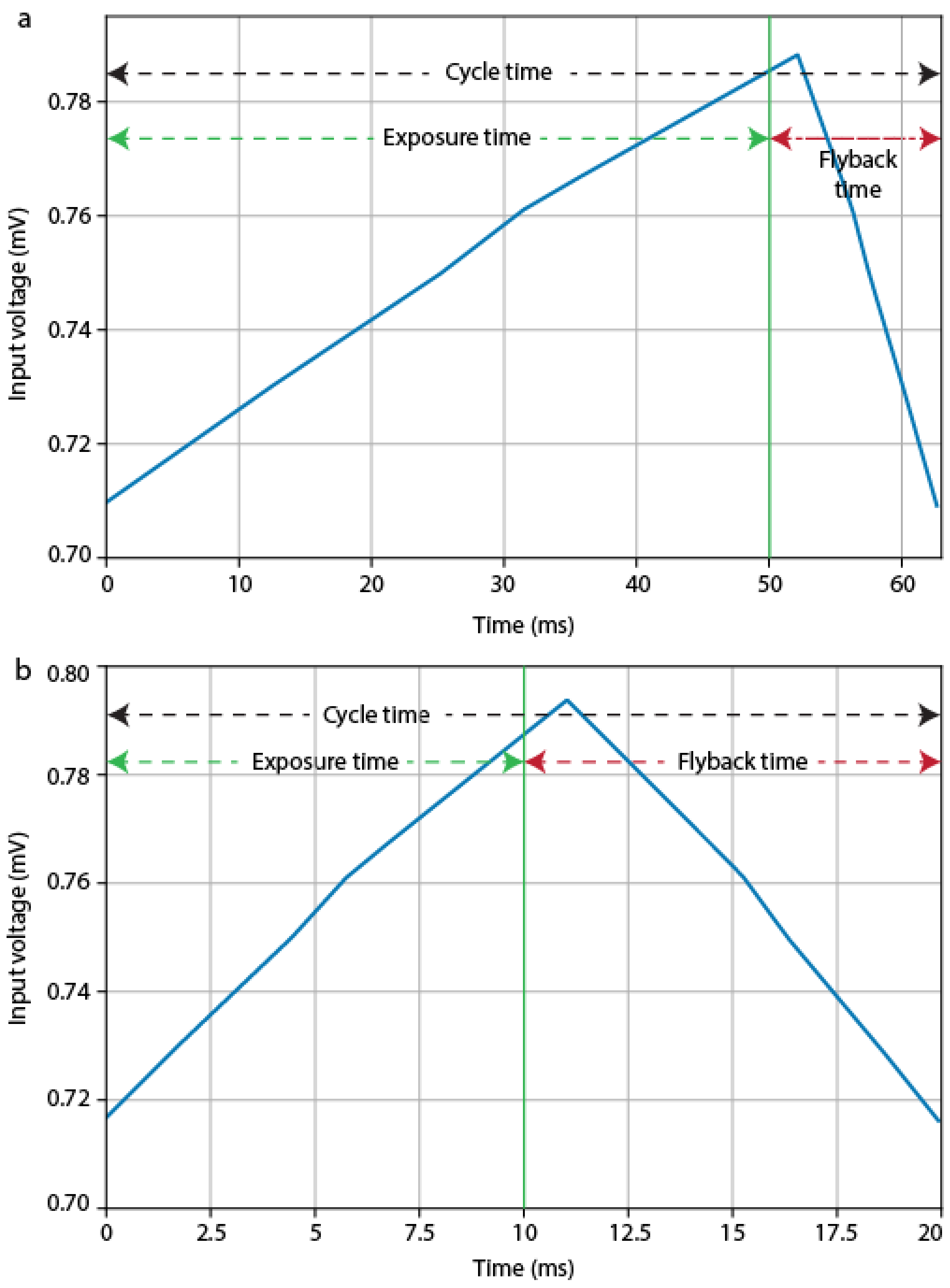


**Supplementary Fig. 20** | **The sawtooth voltage signal for LFA. a,b,** Synchronized TTL triggers for the camera and laser modulation at 50 ms (**a**) and 10 ms (**b**) of camera exposure time. LFA flyback time almost doesn't changes with the decrement of camera exposure time. A signification flyback time causes increment of imaging time though the temporal resolution is equal to the camera exposure time.

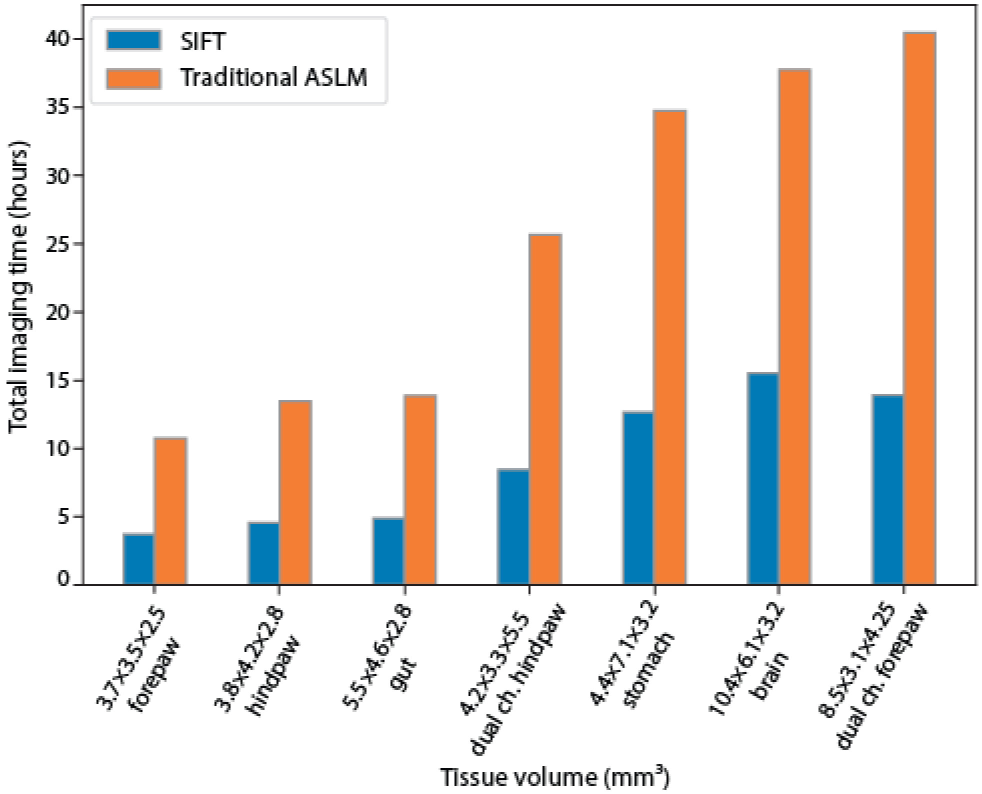


**Supplementary Fig. 21** | **Total imaging time.** Comparison of total imaging time for various tissue specimens having different shapes and volumes. This imaging time is for the equipment used in our experiment (**Supplementary Table 1**). Imaging time may vary depending on the response time of the filter wheel and the stage positional movement.

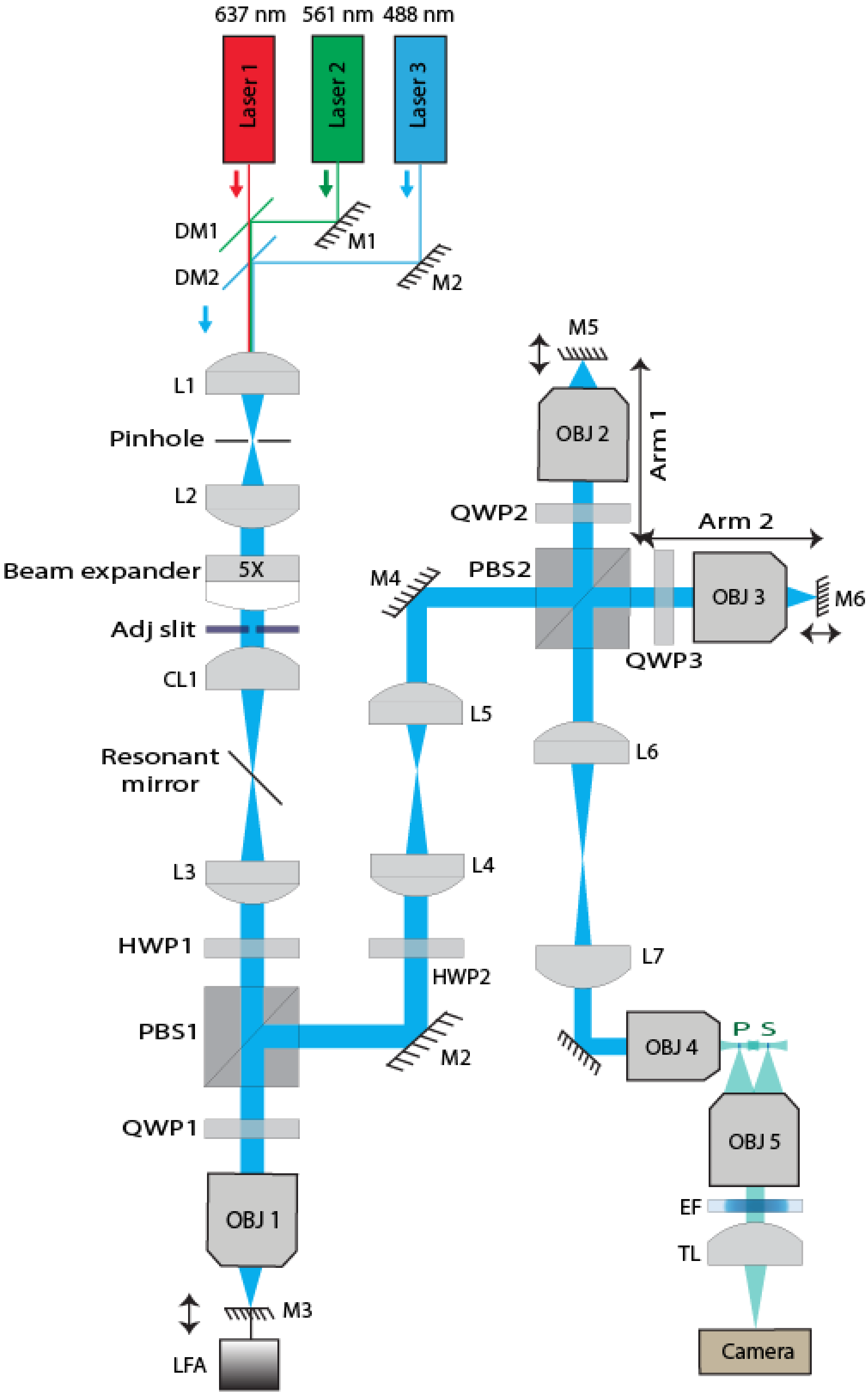


**Supplementary Fig. 22 | Schematic drawing of SIFT.** Corresponding optical components are listed in Supplementary Table 1.

### Supplementary Notes

#### Note 1 - LFA movement

Linear focus actuator (LFA) is a high-performance, small-footprint positioning actuator, specially designed for fast positioning with high precision during short to medium strokes. The voice coil motor is coupled with a high-precision feedback encoder. Low moving mass (50 grams), small step response time (< 3 millisecond) and precise positioning resolution (< 50 nanometers) make LFA useful to various applications such as medical imaging, optical engineering, electronic appliances, semiconductor industry etc.

A tiny mirror is attached in front of the LFA to translate the light-sheet. We found that the LFA movement is not linear with the input voltage over its total travel range[1]. As it is challenging to obtain a precise calibration of the LFA movement with respect to the input voltage, we selected a travel range where the LFA response is linear and covers the range required for imaging the full FOV. With LFA operated in a linear regime, the synchronization procedure becomes easier, where we only need to calibrate the minimum and the maximum voltages for the dual foci to cover the full FOV. And when imaging samples with different refractive indices, we only need to calibrate an offset voltage and keep the difference between the minimum and the maximum voltages the same.

#### Note 2 – Remote focusing technique

A perfect pupil matched remote focusing technique[2–4] is able to correct the aberrations introduced by higher-order polynomials. Though existing techniques such as acoustic lens[5], acoustic LS[6], multi-tiling imaging[7,8], electrotuneable lens (ETL)[9–12] reported to scan LS in the direction of propagation, those techniques introduce additional aberrations.

To achieve perfect imaging (free of aberrations), remote focusing technique demands matching the pupil of the objectives through a 4f imaging system in order to reduce the spherical aberrations. To match the pupils of the two objectives, the magnification ($M$) of the 4f system must satisfy:

$$M = \frac{n_2 F_2 M_1}{n_1 F_1 M_2} \tag{1}$$

where, $M_{1,2}$, $F_{1,2}$ and $n_{1,2}$ represent the magnifications, the focal lengths of the designed tube lenses and the refractive indices of the immersion media for the two objectives. Therefore, a careful selection of the lenses for the 4f system is critical to minimize aberrations. An automatic lens selection software is available for designing the 4f system in remote focusing[14]. Besides reducing aberrations, remote focusing also allows us to image faster. The LFA is capable of sweeping the light-sheet over the entire camera chip (2048 x 2048 pixels) at a speed of 40 frames per second (fps).

#### Note 3 - Building SIFT system

The most promising fact of SIFT is its simple design and ease of installation. We expect the system can be built in two months by an experienced optical engineer. Considering a large tissue, several terabytes of the storage facility and data reading speed are required. The major equipment and parts are listed in **Supplementary Table 1**.

After initial construction, the system is first fine-tuned by generating two 2D focusing beams after the illumination objective and imaging the fluorescein dilution. The two foci are axially swept back-and-forth by LFA. A tight synchronization of the two foci with the camera rolling shutter was achieved by carefully adjusting the range and starting point of the LFA movement along with the flyback time and the cycle time for a specific camera exposure time (**Fig. 2c-2d** and **Supplementary Fig. 20**). A tight synchronization results in a sharp, uniform line over the entire FOV (**Fig. 1**). After synchronization adjustment, a cylindrical lens is added to the illumination path, which generates the light-sheet. For the following alignment we used fluorescent beads embedded in agarose. The rotation and translation of the cylindrical lens were adjusted to achieve uniform bead brightness across the full FOV. The distance between the camera and the detection objective was adjusted to minimize the spherical aberration. A z stack of fluorescent beads was imaged and subsequently, full width half maximums (FWHMs) of the bead images were measured in both the lateral and axial dimensions.

**Supplementary Tables**

**Supplementary Table 1 | Equipment list.** Detail list of materials used to construct SIFT.

| Sl. No. | Description | Item abbreviation | Part number | Part number company | Qty | Remarks |
|---|---|---|---|---|---|---|
| 1 | 4 channel combo OBIS laser | Laser 0 | LX 405-100C | Coherent | 1 | |
| | | Laser 3 | LX 488-50C | | | |
| | | Laser 2 | LX 561-50 | | | |
| | | Laser 1 | LX 637-140C | | | |
| 2 | 427 nm Dichroic beam splitter | DM3 | LM01-427-25 | Semrock | 1 | |
| 3 | 503 nm Dichroic beam splitter | DM2 | LM01-503-25 | Semrock | 1 | |
| 4 | 613 nm Dichroic beam splitter | DM1 | LM01-613-25 | Semrock | 1 | |
| 5 | f = 50 mm, Ø1" achromatic doublet | L1 | AC254-75-A | ThorLabs | 1 | |
| 6 | 30 µm pinhole | Pinhole | P30D | ThorLabs | 1 | |
| 7 | f = 75 mm, Ø1" achromatic doublet | L2 | AC254-50-A | ThorLabs | 1 | |
| 8 | 5X Galilean beam expander | Beam expander | GEB05-A | ThorLabs | 1 | |
| 9 | Adjustable mechanical slit | Adj slit | VA100 | ThorLabs | 1 | |
| 10 | f = 50 mm, Ø1" cylindrical achromat | CL1 | ACY254-50-A | ThorLabs | 1 | |
| 11 | High precision rotation stage | Rot stage | PR01 | ThorLabs | 1 | |
| 12 | Resonant mirror galvanometer | Resonant mirror | CRS 4 kHz | Cambridge Technology | 1 | |
| 13 | 12V DC power supply | 12V DC | A12MT400 | Acopian | 1 | |
| 14 | f = 200 mm, Ø2" achromatic doublet | L3 | AC508-200-A | ThorLabs | 1 | |
| 15 | Half wave plate | HWP | AHWP3 | ThorLabs | 2 | |
| 16 | Polarizing beam splitter | PBS1 | 10FC16PB.7 | Newport | 2 | |
| 17 | Ø1" protected silver mirror | M | PF10-03-P01 | ThorLabs | 5 | |
| 18 | 1" x 1" protected silver mirror | M | PFSQ10-03-P01 | ThorLabs | 3 | |
| 19 | 2" x 2" protected silver mirror | M | PFSQ20-03-P01 | ThorLabs | 2 | |
| 20 | Quarter wave plate | QWP | AQWP3 | ThorLabs | 3 | |
| 21 | Microscope objective (x4, NA 0.28) | OBJ1 | XL Fluor x4 | Olympus Life Sciences | 1 | |
| 22 | Linear focus actuator (LFA) | LFA | LFA-2010 | Equipment Solutions | 1 | |
| 23 | N-BK7 glass piece | GP | 37-005 | Edmund optics | 1 | |
| 24 | Self-Contained XYZ 25 mm translation stage | 3D Tran stage | LX30 | ThorLabs | 2 | |
| 25 | f = 200 mm, Ø2" achromatic doublet | L4, L6 | ACT508-200-A-ML | ThorLabs | 2 | |
| 26 | f = 75 mm, Ø2" achromatic doublet | L5 | AC508-075-A-ML | ThorLabs | 1 | |

| Sl. No. | Description | Item abbreviation | Part number | Part number company | Qty | Remarks |
|---|---|---|---|---|---|---|
| 27 | Microscope objective (x10, NA 0.30) | OBJ2, OBJ3 | | Olympus UMP PlanFl | 2 | |
| 28 | 50 mm travel linear translation stage | Lin Tran stage1 | XR50P | ThorLabs | 2 | |
| 29 | 1/4" travel single axis translation stage | Lin Tran stage2 | MS1S | ThorLabs | 2 | |
| 30 | 2" travel single axis translation stage | Lin Tran stage3 | LT1 | ThorLabs | 1 | |
| 31 | f = 150 mm, Ø2" achromatic doublet | L7 | AC508-150-A-ML | ThorLabs | 1 | |
| 32 | Cleared tissue objective (16.7x/0.4, RI 1.45) | OBJ4 | Special Optics 54-10-12 | Advanced scientific imaging | 2 | |
| 33 | f = 200 mm, Ø2" tube lens | OBJ5 | ITL200-A | ThorLabs | 1 | |
| 34 | Emission filter | EF1 | FF01-525/30-25 | Semrock | 1 | |
| 35 | Emission filter | EF2 | FF01-605/15-25 | Semrock | 1 | |
| 36 | Emission filter | EF3 | BPL01-647R-25 | Semrock | 1 | |
| 37 | 3D motorized stage | Mot stage | Model: MP-285A, PCIe 80 7852R | National Instruments | 1 | |
| 38 | 4 position filter wheel | Wheel | LAMBDA 10-B | Sutter Instrument | 1 | |
| 39 | Digital sCMOS camera | Camera | Orca Flash 4.0, V2 model: C13440-20CU | Hamamatsu Corporation | 1 | |

Qty. quantity

**Supplementary Table 2 | Tissue imaging.** Detail of tissues, imaged using the pipeline.

| Sl. no. | Tissues | Volume imaged ($mm^3$) | No. of tiles | Storage requirement (TB) | Clearing protocol | RI of clearing media |
|---|---|---|---|---|---|---|
| 1 | Mouse forepaw | 3.7 x 3.5 x 2.5 | 685 | 2.22 | Pegasos | 1.56 |
| 2 | Mouse hind paw | 3.8 x 4.2 x 2.8 | 778 | 2.54 | Pegasos | 1.56 |
| 3 | Mouse gut | 5.5 x 4.6 x 2.8 | 878 | 2.84 | Pegasos | 1.56 |
| 4 | Dual channel mouse hind paw | 4.2 x 3.3 x 5.5 | 1,672 | 5.40 | Pegasos | 1.56 |
| 5 | Mouse stomach | 4.4 x 7.1 x 3.2 | 2,171 | 7.03 | Pegasos | 1.56 |
| 7 | Nuclear stained Mouse brain | 10.4 x 6.1 x 3.2 | 2,426 | 7.86 | Pegasos | 1.56 |
| 8 | Dual channel mouse forepaw | 8.5 x 3.1 x 4.25 | 2,588 | 8.38 | Pegasos | 1.56 |
| 9 | Thy-1 GFP neuronal mouse brain | 11.2 x 8.4 x 6.43 | 2,366 | 7.67 | Pegasos | 1.56 |
| 10 | Thy1-YFP-H mouse brain | 17 x 12 x 3.43 | 4,111 | 11.80 | CUBIC-L/R | 1.52 |
| 11 | Mouse colon | 3.7 x 3.7 x 0.23 | 36 | 0.123 | Sca*l*eCUBIC | 1.48 |
| 12 | Zebrafish | 2.8 x 0.7 x 0.62 | 42 | 0.139 | Uncleared | 1.33 |

**Supplementary Table 3 | Multi-immersion imaging.** Detail imaging parameters for multiple tissue immersion media.

| Sl. no. | Refractive Index | Pixel size (μm) | Magnification (x) | FOV ($\mu m^2$) | Remarks |
|---|---|---|---|---|---|
| 1 | Water (~ 1.33) | 0.425 | 15.28x | 870 x 870 | |
| 2 | CLARITY (~ 1.44) | 0.38 | 17.10x | 775 x 775 | |
| 3 | CUBIC-R (~ 1.52) | 0.37 | 17.70x | 750 x 750 | |
| 4 | Tocris (~ 1.53) | 0.365 | 17.80x | 745 x 745 | |
| 5 | PEGASOS (~ 1.56) | 0.36 | 18x | 740 x 740 | |

**Supplementary Table 4 | Cost of tissue imaging.** Comparison between SIFT and the traditional ASLM in terms of the cost of imaging.

| Sl. no. | Tissues | Imaging cost (appr.)[a,b] | |
|---|---|---|---|
| | | SIFT | Traditional ASLM |
| 1 | Mouse forepaw | $ 131 | $ 376 |
| 2 | Mouse hind paw | $ 160 | $ 471 |
| 3 | Mouse gut | $ 171 | $ 485 |
| 4 | Dual channel mouse hind paw | $ 295 | $ 899 |
| 5 | Mouse stomach | $ 443 | $ 1,217 |
| 7 | Nuclear stained mouse brain | $ 542 | $ 1,322 |
| 8 | Dual channel mouse forepaw | $ 487 | $ 1,417 |
| 9 | Mouse colon | $ 17 | $ 50 |
| 10 | Zebrafish | $ 8 | $ 23 |

Appr. Approximated

[a]Included approximated high resolution imaging cost (not included the low resolution and structural evaluation time)

[b]Approximately $35 per hour fluorescence imaging[15–17]
